# The History, Advocacy and Efficacy of Data Management Plans

**DOI:** 10.1101/443499

**Authors:** Nicholas Smale, Kathryn Unsworth, Gareth Denyer, Daniel Barr

## Abstract

Data management plans (DMPs) have increasingly been encouraged as a key component of institutional and funding body policy. Although DMPs necessarily place administrative burden on researchers, proponents claim that DMPs have myriad benefits, including enhanced research data quality, increased rates of data sharing, and institutional planning and compliance benefits.

In this manuscript, we explore the international history of DMPs and describe institutional and funding body DMP policy. We find that economic and societal benefits from presumed increased rates of data sharing was the original driver of mandating DMPs by funding bodies. Today, 86% of UK Research Councils and 63% of US funding bodies require submission of a DMP with funding applications. Given that no major Australian funding bodies require DMP submission, it is of note that 37% of Australian universities have taken the initiative to internally mandate DMPs.

Institutions both within Australia and internationally frequently promote the professional benefits of DMP use, and endorse DMPs as ‘best practice’. We analyse one such typical DMP implementation at a major Australian institution, finding that DMPs have low levels of apparent translational value. Indeed, an extensive literature review suggests there is very limited published systematic evidence that DMP use has any tangible benefit for researchers, institutions or funding bodies.

We are therefore led to question why DMPs have become the go-to tool for research data professionals and advocates of good data practice. By delineating multiple use-cases and highlighting the need for DMPs to be fit for intended purpose, we question the view that a good DMP is necessarily that which encompasses the entire data lifecycle of a project. Finally, we summarise recent developments in the DMP landscape, and note a positive shift towards evidence-based research management through more researcher-centric, educative, and integrated DMP services.

## Introduction

In recent years, significant advocacy and technical enterprise has been directed towards the development and use of data management plans (DMPs) for research. An increase in allocation of library, research office and policy-unit resources towards the promotion of DMPs has led to growing investment in DMP policy and the systems, tools, and educational resources that support it.

We define ‘data management plan’ broadly to be documents that provide researchers with a mechanism for stating how they will manage data associated with at least part of a research project’s data lifecycle. We distinguish ‘data management plan’ from ‘data management plann*ing*’; the former of which is the prepared documentation describing data management practices specific to a research project, whereas the latter is an active process that may or may not involve the preparation of a DMP.

DMPs are frequently promoted across scholarly literature, funding body policy, and institutional communications as a ‘good thing’ for researchers to do, leading to benefits.

We categorise the various claimed benefits as follows:

1. ‘**Professional benefits**’ – the effect of DMP completion on researcher productivity and visibility.
2. ‘**Economic benefits**’ – the effect of DMP completion on increasing the academic and non-academic impact of research, per unit of investment in research funding. These benefits are derived by government, society, and research funding bodies.
3. ‘**Institutional benefits**’ – the usefulness of DMP completion for purposes of institutional planning and compliance.

Economic and professional benefits are promoted as outcomes of DMP use despite that, outside of anecdotes, there has been very limited systematic investigation into the presence or absence of these benefits. The lack of evidence-based inquiry is disconcerting when these untested professional and economic benefits are respectively used to promote the virtues of DMPs to researchers, and used by funding bodies to justify DMP mandates.

The creation, maintenance and oversight of DMPs necessarily impose administrative load on time-poor researchers, and resource-strapped institutions and funding bodies. To justify this, the translation of DMP use to tangible benefits should be demonstrated in practice and provide measurable return on investment.

In this paper, we use a literature search and bibliometrics to trace the DMP from its origins in the 1960s as a bespoke document designed to aid data collection, through to its evolution during the 2000s and 2010s as a planning tool used largely to address funding body expectations for data sharing. We review developments that drove adoption of DMP policies by funding bodies and institutions, and analyse the present policy landscape. We analyse a set of DMPs from a major Australian institution for factors we propose underpin any potential DMP efficacy. We also assess other evidence in the literature that supports and repudiates DMP use, and speculate on the potential for undertaking future investigations to establish whether DMPs have tangible benefits for the communities they were designed for.

## A Brief History of the Data Management Plan

### 1966 to 2000

Results from scholarly literature searches indicate the first evidence of use of data management plans was in1966 (L. W. Ball, 1966; Howell Jr, 1966), where DMPs were used in complex aeronautical and engineering projects. These earliest DMPs were utilised as procedural documents, outlining anticipated research and development activity. These documents were written for project personnel, were freeform in nature, and captured what researchers felt was important to achieve the aims of complex projects.

From the late 1970s to 1980s, DMP use expanded into diverse engineering and scientific disciplines. These DMPs continued to be used as active-stage project management tools, to help complex projects deal with their data management requirements during data collection and/or analysis stages. Exemplar publications include Mason (1975) and Jayroe (1973), both of whom used high throughput instrumentation and complex information systems. Jayroe (1973) was in its entirety a published DMP describing data control in the various steps of computational analysis of a NASA dataset, designed to help other researchers within the field undertake similar projects.

A further example of this active-stage DMP is published as a 1976 conference paper, entitled “Problems of Data Management in a Base Line Study of the Outer Continental Shelf” (Engel & Shaw, 1976). This conference paper discussed past problems with data control in its discipline and described the implementation of data flow control solutions for an offshore drilling study. Similar to Jayroe (1973) this paper reads as an expanded statement of methodology, produced to help others in the discipline. The paper delineates the specific roles of different members of the research team, including project manager, data collectors, and computer centre personnel. The paper describes the details of the flow of data for collection and analysis, reading:

> ‘Procedures are described which have proved a useful plan to insure the success of such a data collection and processing project’ […] ‘Within 10 days of delivery to the computer center, the data from the data collection forms will be punched on computer cards. The data collection forms will be microfiched in accordance with the terms of the contract and the original forms will be filed in the computer center[…] Processing of the data and final entry into storage is done in several steps using programs written in the programming language, PL/1. The data are first recorded in the field using special forms that have been designed to facilitate ease in processing; i.e., cards can be keypunched directly from the forms. The coding form is divided into two sections: the header and the data[…]’ (Engel & Shaw, 1976)

As with most DMPs of this early period, the concept of a ‘data management plan’ is very narrow in meaning. In contrast with modern DMPs, these early papers used DMPs in a manner tailored to solving identified complexity in data acquisition, processing, and immediate (rather than long-term) storage. DMPs were, therefore, used to fulfil a demonstrated short-term need, and were driven by researchers knowledgeable in the requirements of their own projects.

Data management publications of the 1970s to 1980s can typically be seen to address the ‘how’ of the technical implementation of data management processes of a specific study, often at the data collection or analysis stages. Such publications rarely consider the question of ‘why’, in that there is little explicit consideration about the motives driving researchers to use DMPs, other than what we have inferred from what the DMPs contain. One rare exception to this is Mason (1975) in which it was stated:

> ‘In projects of this kind enthusiasm tends to evaporate once the expedition is over and there remain insufficient resources and resolution to extract the full scientific value from the observations. From the outset the Tropical Experiment Board was determined that this should not happen with GATE and accordingly instructed the ISMG to prepare a detailed Data Management Plan and arranged the resources and machinery necessary to conclude the work within 2-3 years.’ (Mason, 1975)

Here, Mason (1975) hints at previous projects lacking the planning and budget for full analysis of the data resulting from their data collection. This suggests that, at least in this case, the DMP was intended to be a means to plan out analyses and budget to ensure that full value would be realised from the collected data. This contrasts with Engel and Shaw (1976) which focussed on controlling the flow of data through collection, processing, and storage systems with particular focus on the roles of staff and the timeframes within which those staff must complete assigned processes.

The two quite distinct uses of Engel and Shaw (1976) and Mason (1975) typify the way in which DMPs were used in this era - that is in an ad hoc manner driven by researchers themselves to achieve project-specific outcomes. Modern ideals held by data professionals around the importance of data sharing, archiving, and reuse, did not have influence on the pragmatic DMPs of this era.

Until the early 2000s, DMPs were utilised in this manner: in limited fields, for projects of great technical complexity, and for limited mid-study data collection and processing purposes, with little cognisance of the concept of the “research data lifecycle” - a term that had yet to appear in the literature.

### 2000 to 2010

In contrast with the organic 20^th^ century development of the DMP, drivers of DMPs in the current century arose from public policy, of which there were two initially distinct but contemporaneous drivers – eResearch and economic policies.

#### eResearch drivers

The early 2000s saw an explosion in global internet traffic and data production, with predictions of doubling events occurring on a near yearly basis (Lyman & Varian, 2003). Hey and Trefethen (2003) and Emmott (2006) argued that this digital revolution was causing fundamental changes to research, where most scientific endeavour shifted to digital processes, and comparatively vast volumes of data were being generated by researchers even outside the traditional domain of ‘big data’. This led to further studies that discussed and speculated more broadly on changes in the nature of research, and the way in which institutions and government should respond to those changes.

One of the earliest and most comprehensive reports on the potential impacts of the then-ongoing shift to the digital age was by Lord and Macdonald (2003); a report commissioned by the Joint Information Systems Committee (JISC; UK). The report discussed the almost complete absence of government and institutional funding for data repositories. It was proposed that the lack of investment in such repositories would rapidly lead to abandonment of and loss of access to huge volumes of data.

A particular concern by Lord and Macdonald (2003) was that most media used at the time, such as CDs, un-replicated hard drives, and tape drives, had lifespans measured in years. Most researchers and institutions would not have made arrangements for longer-term managed storage, effectively leading their proposition that, should their alarm go unheeded, a generation of research data might entirely disappear. Lord and MacDonald (2003) also raised the concern that within science there was at the time a movement for open access to publications, but there was no similar movement for freedom of the data itself. It should be considered that their report was written at a time of remarkably fast obsolescence of computer equipment; much information and communication technology was in the process of rapid standardisation, leaving previous software and hardware to become inaccessible. It was also written at a time when computer deployments within institutions tended to be managed individually with less use of networked or cloud storage, and most researchers unfamiliar with issues of data stability and permanency. On an institutional and commercial level, networked storage was not as well managed, without robust processes for systems redundancy and data backup (Baker et al., 2006; Science and Technology Council, 2007), with the first specifications to standardise data centre integrity only released in 2005 (Standards and Technology Department, TIA, 2005).

To combat these perceived failings of digital technology, Lord and Macdonald (2003) put forward strategies for data management, sharing, and preservation, and advocated for the formulation of data management and curation policies. Their recommendations targeted institutions and government, rather than individual researchers. Lord and Macdonald (2003) stated that it is unlikely that all researchers should require data management and preservation technologies. They also did not describe requirements equivalent to a full data-lifecycle DMP, but rather their recommendations were pragmatic in terms of solving the digital problems identified. Their recommendations focussed on governments and institutions knowing what data exists, and on creating mechanisms to identify and archive data of potential future value.

#### Economic drivers

In early 2004, a group of science and technology ministers from the Organisation for Economic Co-operation and Development (OECD) published a ‘Declaration on Access to Research Data from Public Funding’ (Committee for Scientific and Technological Policy, 2004). The declaration recognised the beneficial impact of open access data, and made a commitment to work towards open access arrangements for publicly funded research data across author countries. The declaration invited the OECD to develop a set of recommendations to be officially endorsed by the OECD council.

This 2004 declaration led to the formation of a working group, with resultant recommendations endorsed by the OECD Council in 2006 (Pilat & Fukasaku, 2007). The recommendations highlighted a perception that OECD countries were suffering low returns on public funding for research because of a near-absence of data reuse. The report discussed the economic benefits of data sharing, but did not consider the professional benefits for researchers. While these recommendations did not directly advocate for use of DMPs, they did make it clear that a responsibility to share data should be placed upon researchers. Most pertinent was their recommendation that “responsibility for the various aspects of data access and management should be established in relevant documents, such as […] grant applications […]”

#### Integration of e-research and economic drivers

A 2005 National Science Board (NSB; USA) report (National Science Board, 2005) drew upon e-research and economic arguments to become the first USA-based report to make the recommendation that a funding body, the National Science Foundation (NSF; USA), require a peer-reviewed DMP with all grant applications. Despite this recommendation, no change in NSF policy is apparent in response to these recommendations.

This same year, of 2005, six major UK research funding bodies funded a consultancy to further interrogate the changing data landscape (Digital Archiving Consultancy, Bioinformatics Research Centre, & National e-Science Centre, 2005). This consultancy report was the first in the UK to recommend that funding bodies should require the submission of DMPs with funding applications. Arguments were largely made from the perspective of the benefits of data sharing and need for controls to be placed on digital data, though there was wider mention of benefits to the researchers themselves, to repositories, to funders, and also reference to the economic arguments of the OECD guidelines. Digital Archiving Consultancy et al. (2005) explicitly took a full data lifecycle approach to DMPs, however at the same time, gave the suggestion that if funders were to implement DMP policies, such policies should be directed towards the funder’s own objectives.

> ‘Our recommendation for data planning recognises that funding institutions and host research organisations will have (or should have) objectives for the data they fund, and strategies and policies for its exploitation. Therefore we believe that specific data plans should be guided by these, and that they should be assessed at or before the point of decisions about funding, and that they should originate from the potential data producer.’ (Digital Archiving Consultancy et al., 2005)

Several UK funding bodies implemented this recommendation, including the Medical Research Council (MRC; UK) in 2006 (Medical Research Council, 2009) and the Wellcome Trust in 2007 (Wellcome Trust, 2008). In both cases, despite the broader data lifecycle view of Digital Archiving Consultancy et al. (2005), the data management policies of the MRC and Wellcome Trust were both, at the time, focused on data sharing to the complete exclusion of other aspects of the data lifecycle. These were the first cases of major funding bodies mandating submission of DMPs with grant applications. A further publication, commissioned by JISC (Lyon, 2007), increased the pressure on funding bodies in the UK to mandate DMPs, following the e-research and economic arguments. Lyon (2007) gave no definition of ‘data management plan’, though it could be inferred from their arguments that they were most concerned with end-of-project data curation and access.

The recommendations made by Digital Archiving Consultancy et al. (2005) and Lyon (2007), that all funding bodies require submission of DMPs with all funding applications, runs in contrast with the more pragmatic intentions of their predecessor Lord and Macdonald (2003), which acknowledged that not all projects should require specific data management considerations. These differences in scope are apparent in DMP implementations by the MRC and Wellcome Trust, where the Wellcome Trust’s 2007 policy required a DMP only in circumstances where a proposal involves a “significant quantity of data that could potentially be shared for added benefit” (Wellcome Trust, 2008).

In a US context, DMP mandates had not yet successfully been introduced at any major funding body until well after those of the UK. In the US, this policy change was spurred by the bringing of e-research and economic drivers together by the Interagency Working Group on Digital Data (IWGDD; US). From 2006 to 2009, the IWGDD, a group representing 27 US Government agencies, prepared a report on data issues in government and society. Recommendation 3 of the final report (IWGDD, 2009) stated that “agencies could consider requiring data management plans for [research] projects that will generate preservation data.” The IWGDD took a broad, full data lifecycle definition of what a DMP should contain.

At this point in the manuscript, we feel it is important to note that while the National Science Board (2005), the Digital Archiving Consultancy et al. (2005), the OECD (Pilat & Fukasaku, 2007), Lyon (2007), and the IWGDD (2009) all strongly advocated that funding bodies require submission of DMPs with funding applications, their recommendations are at best based on workshops and interviews with institutional and governmental stakeholders. Their recommendations and arguments are not backed by quantifiable evidence, nor was significant reference made to researchers themselves. Arguments made in these publications are uniformly devoid of consideration of the professional benefits that DMP use may have directly on the researchers that utilise them, focussing only on e-research drivers, economic drivers, and the nexus of the two.

### 2010 to 2017

#### Funding body data management plan mandates

In response to IWGDD (2009), the National Science Foundation (NSF; USA) announced that all proposals submitted from January 2011 would require submission of a DMP in order to align research practice with the expectation that publicly funded research would be archived and shared (Zacharias, 2010). All projects, regardless of the nature of the project, or the anticipated complexity of the project’s data, require submission of a DMP, even if no research data is to be generated by the project (National Science Foundation, 2010).

In 2013, the Office of Science and Technology Policy (OSTP; USA) disseminated a memo directed to all USA funding agencies to develop a plan to encourage increased public access to any research funded by the federal government (Holdren, 2013), arguing for the economic benefits of data sharing, as well as the principle that the public should have greater access to research it has funded. One of the requirements of these plans is that they would include any necessary policy shifts to mandate DMP use by researchers in receipt of such funds. In total, 18 agencies responded to this, creating and implementing public access plans. In some such plans, including those of the NSF and National Institutes of Health (NIH; USA), agencies reiterated existing DMP policies, however most agencies without DMP mandates introduced them. Resultantly, 62.5% of US funding bodies today require submission of DMPs with funding applications, including most government funding bodies (Table 1).

Different US funding bodies (and, within the NSF, directorates), have different requirements for what elements of the data lifecycle are covered by their DMP expectations. According to a DMP rubric developed by Dietrich, Adamus, Miner, & Steinhart (2012) these DMPs most frequently cover the data archival and sharing stages of a project, and less frequently cover issues such as ethics, intellectual property, and data capture. US funding agencies are in compliance with less than 60% of what Dietrich et al. (2012) considered to be a ‘full data lifecycle DMP’. This is interesting, as funding bodies are mandating DMPs that are consistent with the primary reasons for which funding bodies would want to make DMPs mandatory (that is, the economic benefits believed to arise from data sharing and archiving of end-of-project data). While perhaps not surprising, this is inconsistent with many of the preceding recommendations for DMPs, such as those given by IWGDD (2009) and Digital Archiving Consultancy et al. (2005), supporting a full-data-lifecycle DMP implementation.

In the UK, proliferation of DMP mandates by funding bodies appeared to occur following announcement of NSF requirements. Though as previously described the MRC had a rudimentary DMP policy from 2006, a new wave of DMP mandates by UK funding bodies followed post-2010 (Table 1). The MRC redeveloped its DMP requirements through a JISC-led study involving trialling and receiving feedback about different DMP templates from researchers involved in three MRC-funded studies (Jones, Bicarregui, & Lambert, 2011). The three studies were all epidemiological, and the terms of reference for the investigation was to “trial these templates and give feedback in order to refine them.”

The study was laudable as being one of the only attempts in the literature at gaining feedback about DMPs from researchers themselves, however the study was framed around improving DMP templates, not to determine whether the DMPs themselves hold value. All but one government Research Council in the UK today requires some form of DMP with proposal submission (Digital Curation Centre, n.d.; Table 1).

**Table 1.**
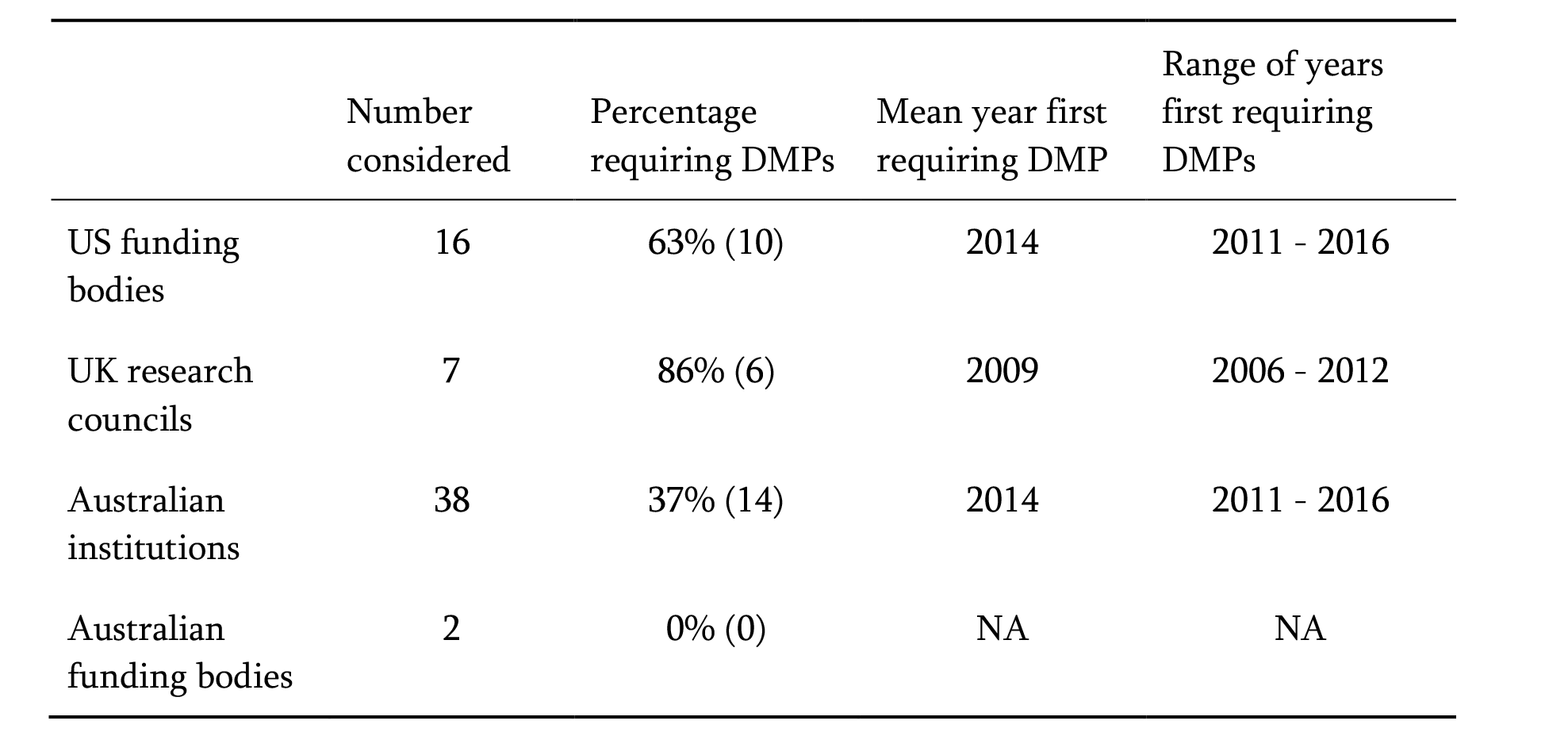
Prevalence of broad DMP mandates from US funding bodies, UK research councils, and Australian institutions. **Methods:** Policies of Australian institutions and international funding bodies were compared through manual compilation of funding bodies and institutions, then using internet searches to source information about the data management policies of these bodies. Year of first DMP requirement was defined as the first year that the body introduced a broad policy requiring that most researchers complete a data management plan. Searches were performed in August 2017. Data available at https://doi.org/10.4225/49/5986bde74f8f5

|  | Number considered | Percentage requiring DMPs | Mean year first requiring DMP | Range of years first requiring DMPs |
| --- | --- | --- | --- | --- |
| US funding bodies | 16 | 63% (10) | 2014 | 2011 - 2016 |
| UK research councils | 7 | 86% (6) | 2009 | 2006 - 2012 |
| Australian institutions | 38 | 37% (14) | 2014 | 2011 - 2016 |
| Australian funding bodies | 2 | 0% (0) | NA | NA |

#### Response of researchers and institutions

In response to funding body-driven DMP mandates, an industry formed to support researchers to comply with these requirements. The first publications to provide general advice and guidance to researchers around the creation of DMPs were published from 2009 (A. Ball, 2010; Donnelly & Jones, 2009; L. Johnston, 2010) following the publications from JISC and the OECD, and after a number of funding bodies had mandated DMP use. Contemporaneous with this is a vast increase in the overall number of DMP-related publications per year (Figure 1). Most DMP advisory publications from 2009 to 2012 eschew reference to direct professional benefits to researchers. Rather, these papers present themselves as a response in support of researchers who must deal with the burden of DMP mandates by funding bodies. DMP use, we infer, has been imposed onto the research community through external forces, rather than through a grassroots effort of researchers and research support staff themselves establishing best practice.

**Figure 1.**
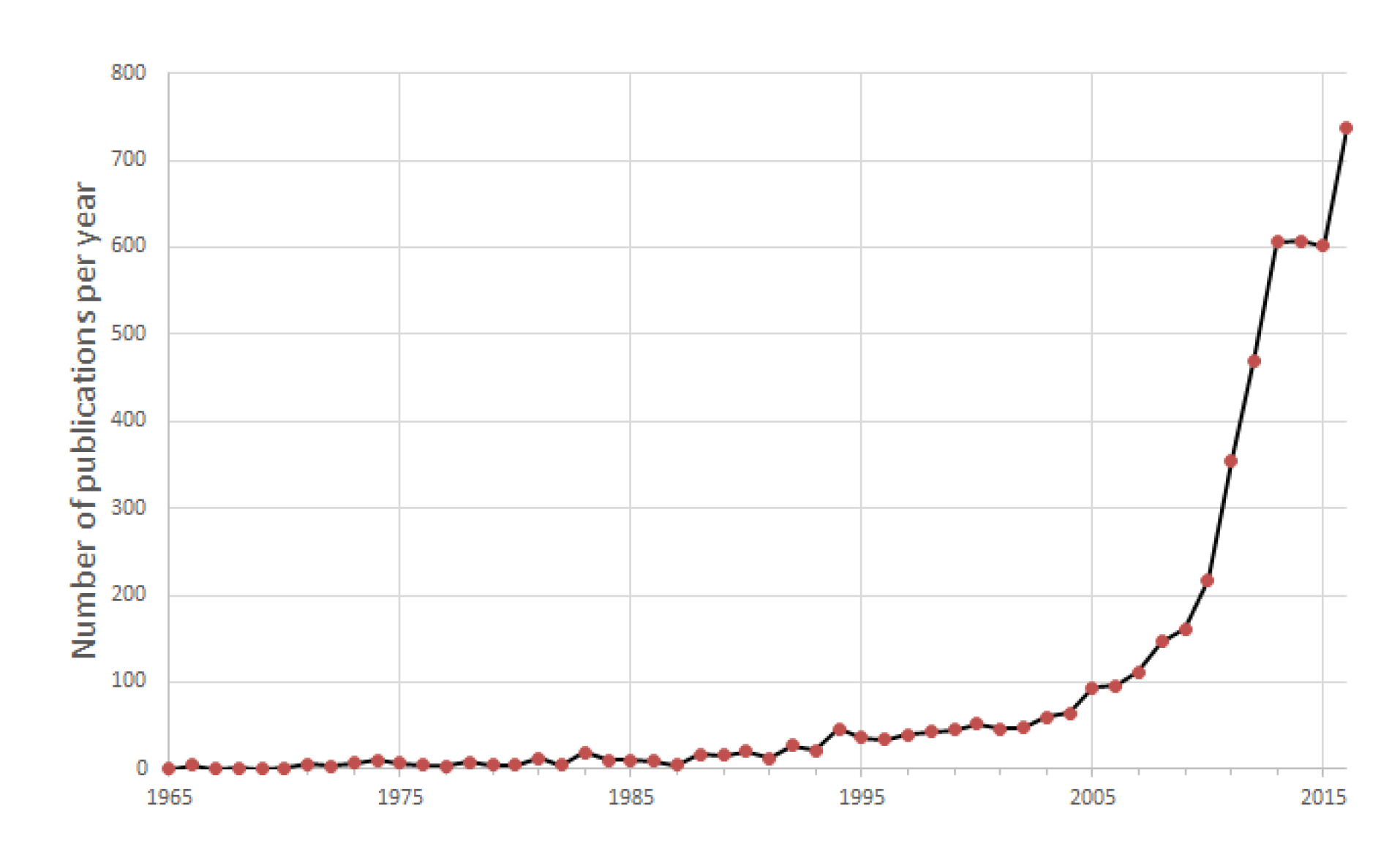
Number of publications per year from 1965 that contain the phrase “data management plan” anywhere in the work. **Methods:** Bibliometric data were collected through searches performed using Google Scholar, querying the phrase, in quotes, “data management plan”. Searches were performed on 8 July 2017.

Announcements by funding bodies of moves towards requiring DMPs with funding applications was a catalyst for university libraries to take on the role of creating and promoting DMP tools and processes (Bishoff & Johnston, 2015; Delserone, 2008). The move towards DMP mandates was contemporaneous with a series of publications urging academic libraries to improve their research data services (Brandt, 2007; Gold, 2007; Salo, 2010; Steinhart et al., 2008). Libraries have capitalised on their strengths in information description, organization and access, deriving benefit from librarian skillsets and positioning themselves as institutional centres of data management practice (Burnette, Williams, & Imker, 2016). Consequently, it is increasingly libraries that have institutionally taken carriage of, and championed, DMPs (Antell, Foote, Turner, & Shults, 2014).

In response to funding body data management plan policies, the Digital Curation Centre (DCC; UK) developed DMPonline in order to assist researchers with completing the DMP sections of funding applications (Simms, Strong, Jones, & Ribeiro, 2016). DMPTool (USA) followed shortly after. These tools were originally developed to help researchers comply with funding body requirements, without imposing an additional administrative workload beyond what the funding body otherwise requires (Sallans & Donnelly, 2012). Though from a UK and USA perspective institutional support structures were at first built to support external DMP pressures placed on researchers by funding bodies, there has since been an expansion of this remit beyond minimal compliance requirements.

In an Australian context, the absence of funder-driven DMP mandates in Australia has led to institution-driven DMP mandates (Table 1). Of institutions in Australia, 37% have DMP mandates in policy. Different models have been adopted: many simply urging researchers to complete static templates, and some linking DMP completion to storage requests and ethics requirements.

## The three benefits of data management plans

### Professional benefit to researchers

There is a common misconception that DMPs were popularised on the basis of professional benefit to researchers. The literature we have reviewed reveals that DMP mandates and resultant popularisation originated from the perception that it was necessary to adapt to the increasingly digital nature of scientific endeavour, as well as a means to encourage the sharing of data for purposes of maximised economic benefit. Through our historical review of DMP development, we have determined that the professional benefits to the researchers who themselves use DMPs have had little more than passing attention paid by those that have sought to mandate DMP use.

Despite this, it is not unusual for researcher-facing publications and websites to promote the professional benefits of DMP use (e.g. Box 1), and for DMP use to be endorsed as ‘best practice’ (e.g. Erway, 2013; Van den Eynden, Corti, Woollard, Bishop, & Horton, 2011). Such claims, however, are only ever accompanied by anecdotal, unreferenced, or unproven justifications. It is our contention that the professional benefits of DMP use have been assumed, and never tested or evidenced. In this section, we assess what evidence there is of DMP use delivering professional benefits to researchers.

#### Box 1

##### Professional benefits to researchers of data management, given within a context of promoting use of data management plans to researchers. This example was taken and anonymised, with permission, from an Australian university.

A data planning process ensures that all aspects of data management are holistically explored at the start of a project. Short-term and long-term aims can be balanced, so that decisions made early in a project do not negatively impact on the ability to find and use the research data in future.

Effective management of data provides researchers with many benefits, including:

- time saved through reduced duplication of effort
- decreased risk of loss, theft or inappropriate use of data
- good research practice ensures the integrity and quality of data
- data can be understood and used now and in the future
- helps researchers find and gain access to data management - expertise and infrastructure offered at the University
- increased researcher profile through data dissemination and re-use.

A data planning process is particularly important in the context of collaborative research projects. Researchers may identify areas of potential difficulty or conflict, and these can be resolved with colleagues and collaborators before they escalate into issues. Clarifying ownership of data, and ensuring early agreement on technical standards and frameworks across institutions, are an important part of establishing trust and ensuring that a project runs smoothly.

There is survey evidence that suggests researchers may have sub-optimal data management skills (Whitmire, Boock, & Sutton, 2015). Researchers report feeling as though they need help with data management, sharing, and archiving (Brandt, 2007). There likely is a service gap in delivering education in data management skills. This gap may be leading researchers to engage in poor data management practices. DMPs are a potential route to assist researchers with data management skills, hence leading to professional benefits. Libraries have opportunistically stepped in to fill this service gap by extending Information Literacy training to include Data Literacy, and by offering research data management support services through partnerships or collaborations with researchers (Corrall, Kennan, & Afzal, 2013; L. R. Johnston & Bishoff, 2015). There may be demand from academic staff for training in relation to data management plans, and such training may be used to address data skills gaps and hence lead to professional benefits to researchers.

In a survey across three Australian universities, 52% of academics stated that they would be interested in training or advice on creating a DMP at the start of a research project (Henty, Weaver, Bradbury, & Porter, 2008). This survey did not explain to respondents what a DMP is, nor state in what form the training or advice would take. The survey also found favourable responses for other forms of data literacy training. Similarly, an American library conducted workshops coaching researchers to produce their own DMPs (L. Johnston, Lafferty, & Petsan, 2012). The authors stated that their workshops were received positively by attendees, however the survey did not assess perceived usefulness or functional outcomes.

Similarly, a study in which library staff educated a small group of interdisciplinary researchers on data management and DMP implementation reported that the process was professionally beneficial to the staff (Burnette et al., 2016). Most notably, a project manager was quoted as saying "I feel like I would have gone crazy” if she hadn’t used a DMP because of the number of protocols and files that were integral to her project.

We do not wholly dismiss anecdotal evidence such as this, however we consider it to be of limited applicability to the majority of researchers. In particular, such evidence could not be generalised to those researchers asked to complete a DMP without associated intensive library support, or those working on less complex projects. Disciplinary differences in attitudes toward data management were captured through a survey by Swan & Brown (2008). The disciplinary perceptions of researchers identified by this survey illustrated these data management differences by contrasting the social sciences view that “(Data management planning) tends not to preoccupy the thoughts of many individual researchers”, with the systems biology view that “Large systems biology teams have a data manager to prepare formal data plans and to manage their implementation.”

Survey evidence does support that researchers have interest in and perceive benefit from DMP-related training. However, such studies to date have failed to directly assess skills or perform longer term follow-up surveys. In particular, these educational interventions have failed to establish any specific benefits of data management training that incorporates DMP use versus data management training that doesn’t incorporate DMP use. The importance of this distinction between training conditions is that there is certainly evidence that researchers derive professional benefits from certain data management practices. For example, the sharing of one’s own research data is associated with an increased citation rate of associated publications (Henry & Fitzpatrick, 2015; Nathan, Genuth, Zinman, & Lachin, 2015; Piwowar, Day, & Fridsma, 2007). Though encouraging sharing of data is one of the main drivers of funding bodies mandating DMPs, evidence of translation from DMP completion to better managed data to more shared data is as yet untested, which raises the question – are there alternate mechanisms besides DMPs that can provide a better means by which data sharing is achieved?

It was argued by Borgman (2012) that though good data practices benefit the researcher, far less documentation is required to maintain data for reuse by the researcher in a future research project than to release those data publicly. It follows that there must be an opportunity cost to maintaining data management at a level higher than that required for use in one’s own immediate research project, where time and money spent on data management is time and money not spent on other research activities. Though we do not necessarily take Borgman’s (2012) view of data sharing, we do accept the broader argument, that time and resources spent training researchers specifically to fill in DMPs might be more efficiently targeted to engage researchers in other forms of data management training.

We acknowledge, from our own experience, that DMP completion can be a catalyst for conversations with institutional staff who can provide help or point researchers to relevant support service providers, such as the Library, IT, Research Office or eResearch specialists. These are services the researcher might not have otherwise known existed had they not completed a DMP. Connecting researchers with these services can enable researchers to gain access to tools and expertise to better manage their research outputs, request storage allocation, discover and deploy collaborative tools, request high performance computing, and access training including statistical analysis, data cleaning, and research data management. A well-designed DMP template can also provide general guidance to researchers on how to avoid downstream issues such as those related to IP, ethics and data publication. In fact, modern DMP tools such as DMPonline and DMPTool devote considerable resources to managing and customising guidance provided for each DMP question.

Despite the above benefits of integrating education and referral services into DMPs, it is our contention that mandatory and poorly thought through DMP templates may be driving researchers to be minimally engaged with the process, applying minimal effort and producing low-quality or insincere DMPs. Given the significant contemporary focus on DMP completion as a means to achieve good data management practices, we are also concerned that the act of completing a DMP, no matter the quality or thought put into the exercise, leads researchers to think that their research data is well managed. Mere DMP completion does not necessitate, predict, or imply good data management practices. Indeed, it is only the start of the data management journey.

There is evidence of a gap between researchers’ ideal and actual data management practices, and this gap is likely impacting data quality and findability, and hence leading to an opportunity cost of professional benefit. We contend however, that there is an incorrect presumption that the training in, or completion of, DMPs organically translates data management theory into best practice data management. We have not been able to find any evidence that DMP use specifically leads to improved data management practices, let alone to downstream professional benefit to researchers.

### Economic benefits to governments and funding bodies

DMPs were mandated by funding bodies largely in response to anticipated benefits from data sharing and reuse. Data sharing is an intuitively beneficial practice with benefits ranging from increased oversight (and, therefore, credibility) of original data and analysis flows, through to the time saved from researchers not having to repeat ‘failed’ (and frequently non-published) experiments.

Various groups have attempted to determine the economic value of data sharing. Economic modelling by Houghton & Gruen (2014) estimated billions of dollars of value is presently unrealised in the Australian research sector due to the low level of data sharing. Piwowar et al. (2011) found that investment in data sharing leads to significantly more publications per dollar spent than direct investment in research. Despite this, Borgman (2012) argues that though much is said of the value of data reuse, there is little information about the potential uses and users of research data. The major reasons why researchers do not release data are often stated to be fear of data-parasitism (i.e. other researchers free-loading on their results) or concerns around making their processed, published results vulnerable to inspection at the raw number level. There is also some evidence that researchers do not release data simply because they cannot imagine who might use them (Mayernik, 2011).

In a study of NSF grant applications at the University of Minnesota, 96% of DMPs made mention of data sharing (Bishoff & Johnston, 2015). However, the vast majority of DMPs in which it was stated that data would be shared, provided responses that did not align with good practice. These included hosting data on personal websites, sharing data only upon request, through conference presentations, and through the non-specific answer of ‘publishing’. These findings suggest that completing a DMP may not be an innately educative experience, with one-size-fits-all funding body DMP requirements not necessarily leading to integrous data sharing practices. Indeed, it has been alleged that it is the immutable vagueness of most funding body data management policies that is in and of itself an impediment to researchers sharing data (Nelson, 2009).

Of DMPs submitted with NSF grant applications at the University of Illinois, there were no significant differences in the proposed data sharing practices of funded and unfunded studies (Mischo, Schlembach, & O’Donnell, 2014). A possible explanation for this is that the NSF peer review process may not select for proposals that are intended to share data. More concerningly, researchers who submit DMPs with funding applications in which they describe how they will share data, frequently do not go on to do so (Van Tuyl & Whitmire, 2016).

Whilst data sharing is likely to improve the quality, reproducibility, leverage and efficiency of research, it is less clear that completion of mandatory DMPs provides the mechanism to ensure that this happens.

### Institutional benefits

DMPs have been promoted as solving institutional compliance requirements, particularly the data documentation, storage, and sharing requirements levied on institutions by funding bodies. Erway (2013), for example, states that DMPs will prompt “…a standardized approach to data management that will ease compliance…”, and discusses the role of institutional compliance offices as stakeholders in DMP design and implementation.

Compliance demands can be based on any of a number of levels, including state or federal legislation, funding body policies, publisher policies, ethics guidelines, and institutional policies. Many of these compliance demands take the form of directives around data retention and sharing. The problem with DMPs being used as a compliance tool in this manner is that compliance is only as effective as the monitoring and remediation that follows. Indeed, it is the experience of the authors of this paper that academics employ a range of strategies to make it appear that they have addressed the requirements of effective data management planning; ranging from extensive copy-pasting from entire grant applications, through overuse of jargon, to perfunctory and provocative assertions. Unless an institution invests in a competent process for the evaluation of and follow-through of DMPs, it is not possible to determine if the DMPs were successful in achieving compliance outcomes. Of course, there is always a proportion of staff that diligently complete a DMP. These good ‘data citizens’ may already engage in good data management and sharing practices, and it may not in fact be this audience that is most in need of support.

There is no doubt that there are enormous potential institutional benefits when all staff practice good data management. The economic and reputational rewards are profound and various: from retention of corporate knowledge through the tracking and reporting of data, integration with workflows dealing with staff and student departures, an evidence trail in the case of research integrity investigations, greater business intelligence around data management and guidance around storage provisioning. Many of these benefits are yet to be realised due to the static and disconnected nature of DMPs, which the move to machine-actionable DMPs may address (Simms, Jones, Mietchen, & Miksa, 2017).

## Analysis of factors relating to DMP efficacy

To date, no rigorous analysis has been published to support, or refute, the relationship between DMP utilisation and the professional, economic or compliance benefits of the previous section. Such an analysis would be a major undertaking, notwithstanding its clear inherent value.

In lieu of any such analysis of DMP efficacy to date, we have sought to perform an analysis of factors we believe are likely pre-requisite to DMPs having any form of significant benefit. These factors relate to the information in DMPs meeting minimal standards of accuracy, completeness, and usability. DMPs that do not achieve basic standards in describing the research data they are nominally about, are unlikely to have the capacity to deliver professional, economic or institutional benefit. On aggregate, such an analysis can provide a system-wide assessment of the capacity of a DMP approach to deliver benefit.

### Methods

A sample was collected from a database containing DMPs from all faculties and disciplines of an established, Group of Eight, Australian university. As is typical of many Australian institutions, this university has internal policies mandating DMP use for all research projects. This DMP mandate is supported by advocacy from library staff, and DMP completion is incentivised through its integration into a compliance checkpoint for users of certain high performance computing resources.

In November 2017, we a random sample of 834 completed DMPs from the university’s DMP database for evaluation across several criteria. DMPs were assessed for (1) detail and quality of information provided about physical and digital data storage (details of classifications given in Table 1).

**Table 1.**
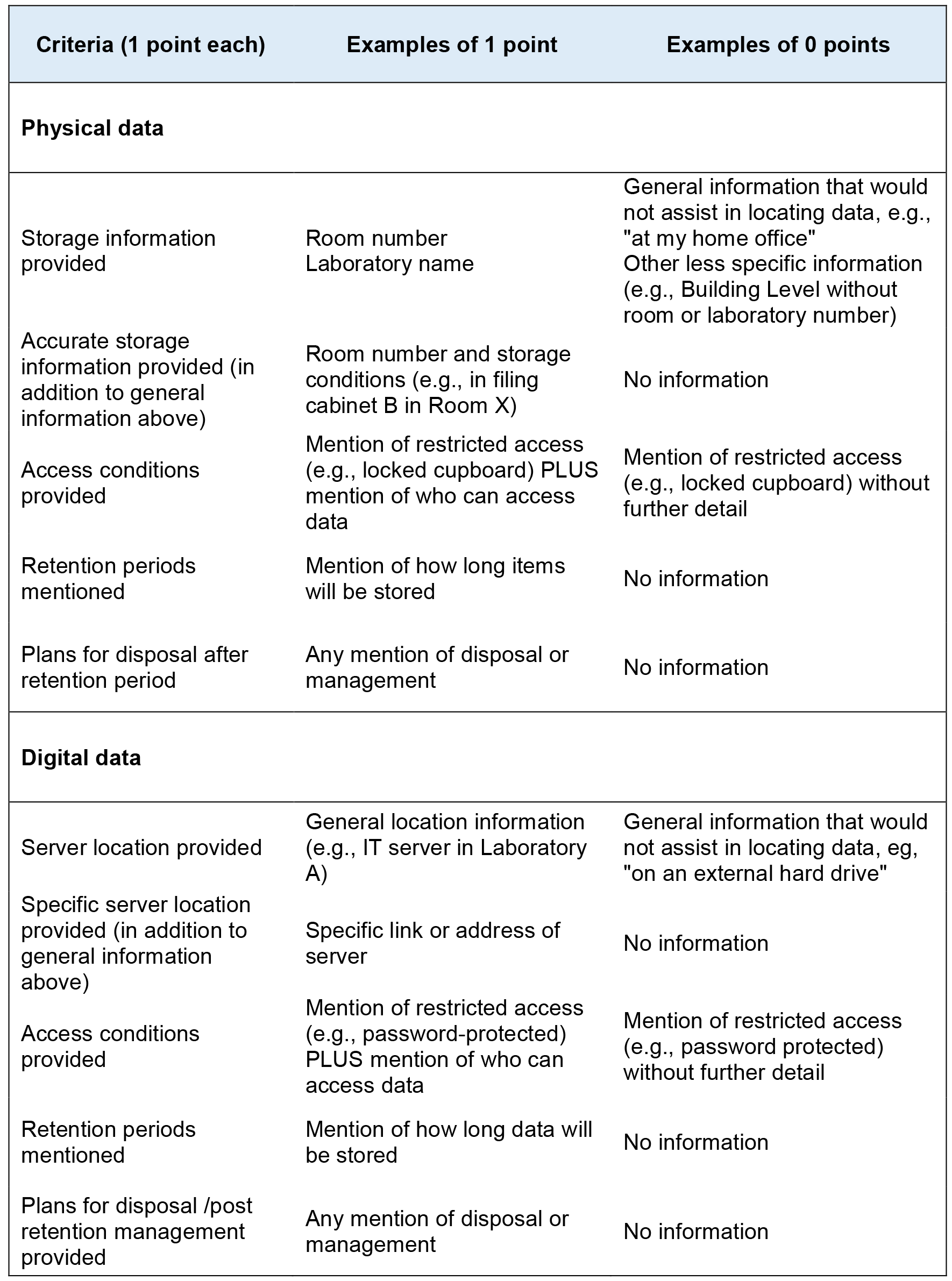
Scoring criteria for physical and digital data storage. Each criterion was awarded 1 or 0 points. Results shown in Table 2.

| Criteria (1 point each) | Examples of 1 point | Examples of 0 points |
| --- | --- | --- |
| **Physical data** |  |  |
| Storage information provided | Room number<br>Laboratory name | General information that would not assist in locating data, e.g., "at my home office"<br>Other less specific information (e.g., Building Level without room or laboratory number) |
| Accurate storage information provided (in addition to general information above) | Room number and storage conditions (e.g., in filing cabinet B in Room X) | No information |
| Access conditions provided | Mention of restricted access (e.g., locked cupboard) PLUS mention of who can access data | Mention of restricted access (e.g., locked cupboard) without further detail |
| Retention periods mentioned | Mention of how long items will be stored | No information |
| Plans for disposal after retention period | Any mention of disposal or management | No information |
| **Digital data** |  |  |
| Server location provided | General location information (e.g., IT server in Laboratory A) | General information that would not assist in locating data, eg, "on an external hard drive" |
| Specific server location provided (in addition to general information above) | Specific link or address of server | No information |
| Access conditions provided | Mention of restricted access (e.g., password-protected) PLUS mention of who can access data | Mention of restricted access (e.g., password protected) without further detail |
| Retention periods mentioned | Mention of how long data will be stored | No information |
| Plans for disposal /post retention management provided | Any mention of disposal or management | No information |

DMPs were also assessed for (2) attitude/effort towards DMP completion and writing quality. Writing quality was assessed according to whether sentences tended to be complete and written to a standard of professional English. Attitude/effort was a subjective judgement as to what extent the DMP was completed with intent, the amount of effort put into answering the questions, and whether sections were left blank. Finally, DMPs were assessed for (3) data type clarity and findability. Data clarity was a binary assessment of how clear it was that the DMP was describing analogue or digital data (or both). Findability was an analysis of whether the DMP contains enough detail to allow a naive third party to locate the digital or physical data (presupposing that any data location details given by the DMP is accurate).

For clarity, we do not suggest these measures as proxy for DMP efficacy. Rather, we propose these measures of DMP completeness, accuracy and usefulness, as a likely dependency of any potential DMP efficacy.

Of 834 DMPs analysed, 21 were excluded because they were identified as being test submissions or submissions intended only to access storage on the University’s Research Data Store; 813 were subsequently included in the analysis.

### Results and discussion

Few DMPs provided specific useful information about the research data nominally being described (Table 2).

**Table 2.** Breakdown of a. Digital scores for DMPs incorporating digital data with or without physical data; b. Physical scores for DMPs incorporating physical data with or without digital data.

| a. Digital Score |  |  |
| --- | --- | --- |
| Score | Number of DMPs | Percentage |
| 0 | 713 | 88% |
| 1 | 81 | 10% |
| 2 | 18 | 2% |
| 3 | 1 | 0% |
| 4 | 0 | 0% |
| 5 | 0 | 0% |
| b. Physical Score |  |  |
| Score | Number of DMPs | Percentage |
| 0 | 497 | 61% |
| 1 | 276 | 34% |
| 2 | 31 | 4% |
| 3 | 9 | 1% |
| 4 | 0 | 0% |
| 5 | 0 | 0% |

In assessment of attitude/effort and writing clarity, the DMPs similarly rated poorly, with only 51% receiving a ‘moderate’ or ‘good’ rating (Table 3). Only 36% wrote in complete sentences, with the remainder having used unprofessional grammar and incomplete sentences.

**Table 3.** Ratings for attitude, writing clarity.

| Attitude/effort |  |  |
| --- | --- | --- |
| Quality | Number of DMPs | Percentage |
| Poor | 399 | 49% |
| Moderate | 359 | 44% |
| Good | 55 | 7% |
| Writing Quality |  |  |
| Quality | Number of DMPs | Percentage |
| Incomplete sentences | 520 | 64% |
| Complete sentences | 293 | 36% |

In many cases, it was difficult to determine even what kind of data was to be produced by the project, and in almost no cases did DMPs describe digital data storage in a way that would be findable (Table 4).

**Table 4.** Ratings for data type clarity, and findability. Scores for ‘physical only’ and ‘digital only’ represent only that one data type is findable, of DMPs that contained both data types.

| **Data type clarity (eg, physical, digital, or both)** |  |  |
| --- | --- | --- |
|  | **Number of DMPs** | **Percentage** |
| Clear | 366 | 45% |
| Unclear | 447 | 55% |
| **Could an external party find the data** |  |  |
|  | **Number of DMPs** | **Percentage** |
| Yes | 24 | 3% |
| No | 513 | 63% |
| Yes - Physical Only | 272 | 33% |
| Yes - Digital Only | 4 | 0% |

Our data overwhelmingly suggest that the vast majority of DMPs within this institutional implementation are likely to be of little or no benefit to the researcher, institution or funding body, given that they do not appear to describe a plan for data management. This particular implementation represents almost 2000 researchers over the multi-year lifespan of the system having been compulsorily made to partake in an administratively burdensome exercise, with no identifiable tangible or intangible benefit. We highlight that it should be the responsibility of DMP system owners to identify and take measures to correct trends of apparent inefficacy.

We do not intend to generalise these findings to all DMP implementations, however we do find these results concerning.

## Current thinking and future directions

In this paper, we have identified several issues with the current use of DMPs. As argued here, much of the advocacy around DMPs assumes professional benefit to researchers. However, there is no apparent evidence-base upon which to support the argument that researchers gain professional benefit through the act of completing a DMP, nor is there evidence as to whether DMPs are an effective way of approaching the data management skills gap. Our analysis of DMPs from one Australian institution suggests that DMPs may in fact be ineffective, for the basic reason that they generally do not actually describe research data management to any acceptable level of quality or detail.

In recent history, DMPs have been promulgated by funding agencies on the basis of encouraging data sharing. Despite this, even when researchers use a DMP to identify how and where they will share their data, most researchers do not follow through with their own plans. Indeed, despite interrogation of the literature and extensive inquiries made to members of the data management community, we have been unable to identify any systematic evidence that supports the view that DMP completion has any innate benefit to any party. Simultaneously, the inconsistency in application of the term ‘data management plan’ amongst funders, institutions and researchers is adding to the current lack of integration of effective data management planning.

We suggest using outcome-focused approaches to align DMP use with specific goals. Delsersone (2008) differentiates between diverse forms of data management: that which is driven by a particular discipline’s standards, data management driven by the concerns of an individual researcher, data management planning at the behest of a specific funding agency, and institutional data management planning. DMPs have become a tool for addressing the requirements of multiple stakeholders. Perhaps these different drivers need to be better delineated, and the interests of each identified, with an evidence base to support the approach taken.

Work is required in identifying DMPs, or alternative mechanisms, that are “fit for purpose”, where that purpose is explicit, for example that which meets requirements from funders, or business intelligence gathering by institutions, or as a change management tool for behaviour modification, or project management planning for researchers. This becomes a non-trivial task when multiple purposes are being addressed at once.

### Use by funding bodies to encourage data sharing

Over the past decade, funding bodies have mandated DMPs as a ‘soft touch’ approach to directing researchers to share their data. However, current DMPs may be counterproductive in that they saddle researchers with unnecessary administrative burden, and encourage meeting minimum requirements. If funding bodies pursue their current data sharing imperative, then the ideal DMPs should: 1) focus on the mechanics of data sharing at the latter end of the data lifecycle; 2) be publicly accessible so that users can find how to access past projects’ data; 3) ensure grant applications are assessed by a grant review panel with terms of reference and a weighting that strongly selects for data sharing projects where appropriate; and, 4) provide compliance mechanisms that are visible, explicit and demonstrate accountability.

It may be that DMPs are being used to achieve aims that could be better accomplished by other instruments. A variant on the DMP approach is the use of data sharing plans (DSPs). DSPs are intended to more explicitly focus on how the applicant would comply with the funding body’s data sharing policy, focussing on the latter end of the data lifecycle. The nomenclature of ‘DSP’ also dispels some of the confusion and inconsistency around what the objective of the task is. The largest funding body to implement such a system is the National Institutes of Health (NIH; US), which has since 2003 (NIH, 2003) required DSPs with project applications.

The NSF’s directorates, and indeed most funding bodies, have DMP requirements that are consistent with, though in excess of, a DSP (Dietrich et al., 2012). These require detail not relevant to their stated aims of encouraging data sharing. The practice of requiring researchers to complete forms that cover potentially all aspects of research data management should be reviewed, with a refocus on asking researchers to explain how, where, and when they will share their data, or give researchers the opportunity to justify why they are unable to do so. Where researchers identify that they will share their data, detail should be included about specific repositories. Because publications add context and value to data (Borgman, 2012; Pepe, Mayernik, Borgman, & Van de Sompel, 2010), applicants should also make clear the links between publications and the associated data, and preferably any open data mandate should be matched with an open access publication mandate.

Casting doubt on the efficacy of DMP/DSPs as a mechanism for data sharing is the apparent lack of compliance by researchers, who tend not to follow the data sharing protocols outlined in the DMPs that accompany their own funding applications (Van Tuyl & Whitmire, 2016). The existing honour system, where researchers complete DMPs/DSPs, then are expected to comply post-hoc, does not appear to be effective. Even blanket data sharing policies by publishers are proving difficult to enforce. In one study of datasets associated with PlosOne papers, the researchers were only able to acquire the datasets underlying one of ten publications - despite the authors of all ten papers having agreed at the time of publication to share their data (Savage & Vickers, 2009). Wiley’s 2016 Data Sharing Survey updates figures from their 2014 survey show an increase from 52% to 69% of researchers being prepared to share data, but only 41% report sharing data via data repositories (Wiley, 2016). Attempts at creating cultural change through funder DMP mandates and journal data sharing policies, although laudable, likely remain ineffective.

In contrast with most funding bodies, the Australian Antarctic Science program takes into account the researcher’s previous history of data sharing when assessing proposals for funding, leading to a demonstrable positive effect on researcher data management practices (Finney, 2014).

If the economic value intrinsic to data sharing is commensurate with dollars estimated in various reports (e.g. Haughton & Gruen (2014) calculating up to $4.9 billion of unrealised economic benefit within Australia), then the amount of money lost through researchers not complying with data sharing conditions is probably much greater than that lost through any other form of grant funds misdirection or fraud. Viewed from this perspective, it would follow that data sharing representations made by researchers should be taken much more seriously, and instilled in researchers that not complying with such representations could be considered a breach of research integrity. Data sharing representations made by researchers in their funding proposals, whether through DMPs, DSPs, or otherwise, could be more actievly audited for compliance.

A non-DMP approach with potential for satisfying the data-sharing use-case is to employ institutional mandates that require data sharing. Institutional open access mandates are effective at increasing the proportion of publications for which open access arrangements are made, and there is a correlation between the strength of the institutional mandate and the total proportion of researchers in compliance with that mandate (Gargouri, Larivière, Gingras, Carr, & Harnad, 2012). When staff performance evaluations are linked to compliance with a mandate, policy is more likely to be followed. An additional benefit of such a system may be that linking staff performance metrics with data outputs could help drive a cultural shift towards establishing data as a valued ‘first class’ research output.

### Use by institutions to change researcher behaviour

When the desired purpose of a DMP is to drive researcher behavioural change in data management practices, DMP completion must be seen as a value-add for researchers and include an educative focus. Ideally the DMP system used would: 1) allow for iterative interaction for the purpose of review and updating; 2) focus on the data collection, processing and analysis methodologies/methods relevant to the specific field of research; 3) be easily integrated into researcher workflows; and, 4) be scaffolded through DMP training (face-to-face or online) and other associated resources and materials that form part of a DMP system, potentially consisting of:

- A training manual/online modules
- An institutional storage options chart
- A referral map of all research support services across the institution
- A DMP self-assessment rubric.

If completion of a DMP is to be considered a tenet of good data management practice and a behavioural change management tool for researchers, then there should be evidence of its efficacy. Part of this evidence seeking would need to explore factors that influence adoption of good practice in data management and test claims of professional benefits presently assumed to be derived from DMP use. Such analyses could attempt to correlate DMP use with researchers gaining higher profiles, saving time, or increasing their productivity. Such evidence would be self-reinforcing, by providing the evidence-base that could be used to convince researchers that these benefits are tangible.

Further research could also be undertaken to explore what impact DMPs have on grant proposal success. For specific funding bodies that require DMPs, does DMP quality actually have a significant impact on grant success? If it could be proven that poor DMPs lead to rejected grant applications, then completing a DMP to a higher standard could lead to greater grant success and be seen as a professional benefit to researchers, albeit one that is artificially produced through funder mandates. Indeed, a 2012 survey found that 87% of researchers intending to submit NSF grant applications felt that they would benefit from advice or help with complying with NSF DMP requirements (Steinhart, Chen, Arguillas, Dietrich, & Kramer, 2012). A similar 2013 study utilising a survey and interviews found that most researchers expressed a great deal of uncertainty around data management, and around half would like further assistance with developing a data management plan (Rolando, Doty, Hagenmaier, Valk, & Parham, 2013).

Discussions at an international level are beginning to address aspects of DMP quality against specific criteria using assessment rubrics (Parham, Carlson, Hswe, Westra, & Whitmire, 2016), but how this scales given limited human resources to undertake such assessments remains an open question.

### Use as an institutional business intelligence and systems integration tool

If being adopted as a component of institutional business intelligence gathering and systems integration, then DMPs and existing institutional systems should be connected where possible to alleviate duplicating the collection of the same content and enable easier integration with researchers’ workflows. These DMPs should be:

- Online and enable future reviews/updates
- Integrated with business systems relating to:
  ○ proposal preparation, submission, and tracking
  ○ storage services, providing a trigger for storage allocation requests
  ○ ethics approval processes
  ○ the institutional metadata repository to enable the creation of pre-publication records, application of an appropriate licence, reservation of a DOI and a data citation
- Able to trigger alerts to the institution’s Industry Engagement/Commercialisation unit
- Inclusive of a retention/disposal flag
- Exportable as PDFs for funders, publishers and institutional reporting purposes.

The benefits of DMP use for business intelligence gathering and systems integration may be demonstrated through aggregate DMP data being an input into other decision making processes at the institution. DMP use might allow institutions to plan for the digital and physical storage space, data management tools, and staff and services to support the predicted needs of researchers, though with a need to temper these predictions with the possibility that researchers may not always be capable of accurately relaying or understanding these needs. Use of DMPs purely as a mechanism for business intelligence gathering and/or systems integration would need to be weighed against the needs for educational and behavioural change within research communities, factoring into decision making the need for training as part of a research support program.

### Use by researchers for project management

Looking back to the 1960s and 70s, DMPs filled the needs of researchers or research teams in guiding how they managed the data coming out of complex projects. This style of planning enabled researchers to take their own approaches to managing data based on their specific circumstances. Data was not necessarily separated out as a special case, but formed part of overall research project planning as an intrinsic and underpinning component of the research itself.

Taking such a researcher/research-centric approach means that discipline norms, methodologies and methods related to data collection, capture, processing and analysis can be explicitly captured and not lost due to the generic fog of the contemporary DMP template. Therefore, institutions should provide services to help researchers project manage their research and research data. These could be responsive to prompting detail about the parts of the data lifecycle that the researcher is by necessity interested in.

The effectiveness of DMPs in project management would not particularly need to be evidenced as it would be up to individual researchers to opportunistically use DMPs in this manner.

## Conclusion

DMPs began life as a practical document used by researchers for mid-project data collection and analysis. This narrow and researcher-led use case gave way to concerns about the changing and increasingly digital nature of science, in addition to concerns about lost value through data not being shared. Funding bodies and institutions have increasingly acted to make DMP use mandatory, and to use DMPs as a proxy for evidence that data management planning has taken place. Nevertheless, there is a paucity of evidence that any real professional, economic or institutional benefit is gained from the act of completing a DMP. Our own analysis of a sample of 834 DMPs from a typical Australian institutional DMP implementation found little intuitive opportunity for the realisation of proposed benefits. In the worst case scenario, DMPs may add only administrative load to researchers.

We do not seek to condemn institutional responses to the changing DMP requirements of funding bodies. Researchers who intend to submit, for example, NSF grant applications do feel that they need advice with producing adequate DMPs, justifying the institutional provision of certain DMP services. We similarly do not intend to criticise funding bodies that have often been led by government policy in mandating DMPs.

It may be that the questions asked by DMPs may not have previously been considered by the majority of researchers and so may play some part in enacting a cultural shift in research towards consideration of data lifecycle issues. Nevertheless, the present publication came about as part of preparation of a research data management training course, due to a desire to communicate to researchers the importance of good data management practices. We sought evidence that DMPs are beneficial to researchers, in the hope that this would cause attendees to adopt DMPs into their own projects. An extensive literature review and informal enquiries have found scant evidence that DMP use is to the benefit of any party.

Though the UK and USA have over the past decade followed a model of funding body mandates for DMPs, we suggest that in the absence of evidence, these additional administrative burdens offer little or no tangible efficacy in affecting research practice. This should be a point of concern for data management professionals and policy-makers.

We acknowledge exciting developments presently occurring across the DMP landscape, towards more researcher-centric, educative, and integrated DMP services. However, we put forward that more consideration needs to be given towards the aims and objectives (the ‘use case’) of DMPs, and that DMP implementation needs to be outcome-driven in a manner consistent with those use-cases. The efficacy of DMPs specific to their proposed use case should be demonstrable and measurable to ensure mandates are defensible and have a net positive impact on the research endeavour.

## Acknowledgements

The authors wish to acknowledge Peter Neish, Research Data Curator at the University of Melbourne, for providing extensive insight and advice in preparing this manuscript. We acknowledge Dr Elise Magatova for providing advice on data analysis.

Additionally, we thank Dr Richard Ferrers, Research Data Analyst at the Australian Research Data Commons, for invaluable suggestions made to an earlier version of this manuscript. Finally, we dearly thank members of the Australian and international data management communities who have generously and patiently assisted us with enquiries in relation to this manuscript.

